# Distributed Sampling-based Bayesian Inference in Coupled Neural Circuits

**DOI:** 10.1101/2020.07.20.212126

**Authors:** Wen-Hao Zhang, Tai Sing Lee, Brent Doiron, Si Wu

## Abstract

The brain performs probabilistic inference to interpret the external world, but the underlying neuronal mechanisms remain not well understood. The stimulus structure of natural scenes exists in a high-dimensional feature space, and how the brain represents and infers the joint posterior distribution in this rich, combinatorial space is a challenging problem. There is added difficulty when considering the neuronal mechanics of this representation, since many of these features are computed in parallel by distributed neural circuits. Here, we present a novel solution to this problem. We study continuous attractor neural networks (CANNs), each representing and inferring a stimulus attribute, where attractor coupling supports sampling-based inference on the multivariate posterior of the high-dimensional stimulus features. Using perturbative analysis, we show that the dynamics of coupled CANNs realizes Langevin sampling on the stimulus feature manifold embedded in neural population responses. In our framework, feedforward inputs convey the likelihood, reciprocal connections encode the stimulus correlational priors, and the internal Poisson variability of the neurons generate the correct random walks for sampling. Our model achieves high-dimensional joint probability representation and Bayesian inference in a distributed manner, where each attractor network infers the marginal posterior of the corresponding stimulus feature. The stimulus feature can be read out simply with a linear decoder based only on local activities of each network. Simulation experiments confirm our theoretical analysis. The study provides insight into the fundamental neural mechanisms for realizing efficient high-dimensional probabilistic inference.

## 1 Introduction

The theory that the brain performs probabilistic inference to interpret the external world [1–3] has been supported by extensive human behavioral [4–6] and animal neurophysiological studies [7, 8]. Yet, exactly how neural circuits in the brain realize Bayesian inference remains poorly understood. To address this question, we need to answer how probabilistic information is represented in neural responses and what inference algorithms are adopted by neural circuits. An added difficulty is that signals in the world are high-dimensional. The brain often extracts and represents multiple stimulus variables in a parallel, distributed, and hierarchical fashion using separate neural circuits. Examples include: integrating multisensory cues [8], resolving the consistency of local percepts (e.g. nose and eyes) and global percepts (e.g. face) in a hierarchy [1, 9], and grouping orientation edge neurons into contour [10, 11]. The brain needs to represent and infer the joint posterior distribution of all these variables in different locations, different levels, or even different sensory systems. This fundamental challenge must be addressed in the context of other properties and constraints of Bayesian inference in the brain.

The neural mechanics of inference has been an active research area, with a number of existing computational models. A popular proposal is that of probabilistic population coding (PPC), whereby a deterministic network integrates feedforward Poissonian inputs to parametrically infer and represent the posterior of a stimulus feature [12]. This model was later extended to process high-dimensional stimulus features by coupling networks together, where the coupling encodes the correlation prior between stimulus features [13]. Such a model scales linearly, rather than combinatorially, with the number of stimulus features. This allows each feature’s marginal distribution to be computed and readout from each network. Despite its conceptual appeal, the interaction in such a coupled PPC network is highly nonlinear, requiring a complex circuit with multiplicative and divisive operations. An alternative paradigm to represent the joint posterior of multiple variables is using non-parametric sampling (e.g., [14–20]), which can be mediated by linear stochastic dynamics (e.g., [16]). However, these studies typically consider sampling the posterior of neuronal responses, without specifying how stimulus features embedded in neuronal responses are sampled (e.g., [14–16]), or they considered sampling stimulus features but did not specify a concrete neural circuit model to enact the sampling [18, 21]. Moreover, when sampling occurs in the very high-dimensional neural response space, involving neurons in different hypercolumns, visual areas, and even different sensory systems, the sampling timescale becomes a serious obstacle [16, 22], which must be overcome to produce perception in a biologically realistic time frames.

In this study, we propose a novel model to represent and infer the posterior of high-dimensional stimulus features. Our model combines the strengths of PPC and sampling-based codes in a unified framework. It decomposes the high dimensional parameter space into distinct variables, represented individually by distinct continuous attractor neural networks (CANNs). In our model feedforward inputs convey the likelihood via PPC, excitatory reciprocal connections between the attractors encode the stimulus correlational priors, and the internal Poisson variability of the neurons generate the correct variability for sampling on the stimulus feature space. We analytically show that the dynamics of coupled CANNs in fact realizes Langevin sampling on the stimulus feature manifold embedded in neural population responses. The model achieves sampling-based Bayesian inference in a distributed attractor network, each of which infers the marginal posterior of the corresponding stimulus feature, and its local activities allow readout of the stimulus feature using a linear decoder. Simulation results confirmed our theoretical analysis of the model.

## 2 The generative model and sampling-based inference

### 2.1 The probabilistic generative model

We consider a linear Gaussian generative model (Fig. 1A), in which the observed stimulus features 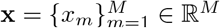 are independently generated by the external latent features 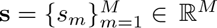. These features can be any stimulus features that are extracted by the cortex, such as line orientation, moving direction etc. The likelihood function *p*(**x**|**s**) and the prior *p*(**s**) of the generative model are

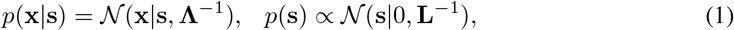

where 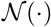 denotes a Gaussian distribution. **Λ** is the precision matrix (the inverse of the covariance matrix) of the likelihood function, which is assumed to be diagonal, i.e., **Λ** = diag(Λ_1_, Λ_2_, ···, Λ_*M*_). This implies that each observed feature *x*_*m*_ is independently generated by *s*_*m*_, giving 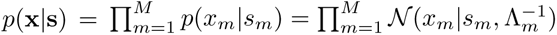.

**Figure 1:**
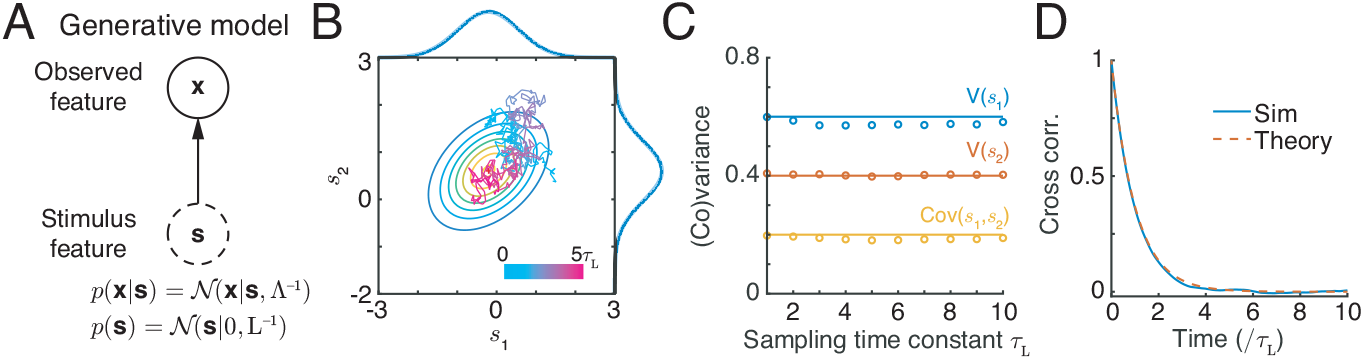
A probabilistic generative model and its inference by Langevin sampling. (A) A linear Gaussian generative model, where **s** and **x** are the latent and observed stimulus features, respectively. **Λ** and **L** are precision matrices of the likelihood function and the prior, respectively. (B) Langevin sampling for approximating the posterior of two-dimensional features. Solid ellipses: the posterior. Colors of the trajectory indicate the time elapsed. Marginal plot: marginal posterior (empirical: solid line; theory: shaded line). (C) The equilibrium variance of samples vs. the sampling time constant *τ*_*L*_ (empirical: circle; theory: solid line). (D) The temporal correlation of sampling over time.

The precision matrix **L** of the prior *p*(**s**) is a generalized Laplacian matrix, with L_*mm*_ = −∑_*n*_ L_*mn*_ and L_*mn*_ = L_*nm*_ ≤ 0 (*m* ≠ *n*). For example, in the case of two-dimensional features (*M* = 2), the prior *p*(**s**) = *p*(*s*_1_, *s*_2_) is written as,

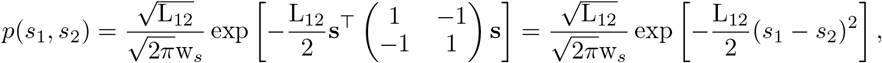

where w_*s*_ is the width of stimulus feature space, e.g., w_*s*_ = 2*π* if *s* is direction of stimulus motion. Priors with a Laplacian precision matrix have been *implicitly* considered in the modelling studies for multisensory cue integration [23–25], and it quantifies the co-occurrence of stimulus features, e.g., L_12_ characterizes the correlation between *s*_1_ and *s*_2_. Notably, according to Eq. 1, the marginal prior of each stimulus feature is uniform, i.e., *p*(*s*_*m*_) = 1/w_*s*_, a property coming from that the determinant of the Laplacian matrix is zero, i.e., |**L**| = 0.

According to Bayes’ theorem, the posterior distribution of the latent variables **s** given the observed features **x** are calculated by inverting the generative model (Eq. 1),

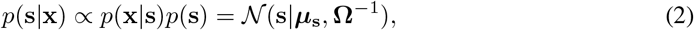

which is a multivariate Gaussian distribution, with the precision matrix **Ω** and mean ***μ***_**s**_ given by,

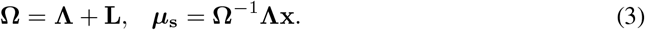

### 2.2 Langevin sampling for approximate inference

Numerous psychophysical studies suggest that the brain infers the external world via Bayesian inference (e.g., [2, 3, 26]), but it remains unresolved how the inference is implemented by cortical circuits in the brain. Here, we propose that the brain employs a sampling strategy to approximate the inference of stimulus features, and later we demonstrate that this is feasible in a biologically plausible neural circuit. The sampling strategy we consider is Langevin sampling, which is a Markov chain Monte Carlo (MCMC) algorithm widely used to numerically approximate the posterior [16, 27, 28]. The dynamics of Langevin sampling performs stochastic gradient ascent on the manifold of the log-posterior of stimulus features, which is written as,

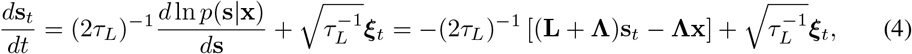

where *τ*_*L*_ is the time constant of sampling. ***ξ***_*t*_ are multivariate independent Gaussian-white noises, satisfying 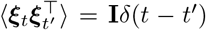, with **I** the identity matrix and *δ*(*t* − *t*′) the Dirac delta function, and they induce fluctuations of **s**_*t*_ necessary for sampling. It can be checked that the equilibrium distribution of **s**_*t*_ has the same form as the posterior (Eqs. 2-3), since its equilibrium mean and covariance satisfy (see details in Supplementary Information (SI.) Sec. 2),

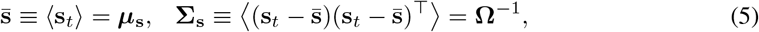

where 〈·〉 denotes averaging over trials. Thus, in the dynamics of Langevin sampling, the instant value of **s**_*t*_ can be regarded as a sample from the posterior specified by Eqs. (2-3). Fig. 1B shows an example trajectory of Langevin sampling of two-dimensional stimulus features **s**_*t*_ over time, demonstrating that the sampled **s** indeed satisfies the posterior as expected. The sampling time constant *τ*_*L*_ doesn’t affect the equilibrium covariance (Eq. 5), but only affects the temporal correlation of sampling (i.e., the normalized cross-correlation of samples at time *t*_1_ and *t*_2_, which is given by 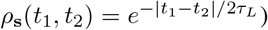, and these two theoretical predictions are also confirmed by simulations (Fig. 1C-D).

## 3 Distributed sampling-based inference in coupled attractor networks

### 3.1 Coupled continuous attractor neural networks

We explore how a canonical neural circuit model implements Langevin sampling to approximate the posterior of high-dimensional stimulus features. The model we consider is composed of *M* reciprocally connected continuous attractor neural networks (CANNs), and they interact with each other to achieve inference in a distributed manner, such that each network *m* infers the *marginal* posterior of one stimulus feature *p*(*s*_*m*_|**x**). For simplicity, we consider that each network *m* has the same number of *N* neurons with preferred stimulus values 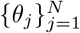, with *θ*_*j*_ being the preference of the *j*-th neuron. We take 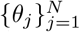 to uniformly cover the feature space of *s*_*m*_, where *θ* ∈ (−*π*, *π*] satisfying the periodic boundary condition.

CANNs are a canonical neural circuit model widely used to elucidate the network mechanism underlying many brain functions (see e.g., [29–31]). In the continuum limit (*θ*_*j*_ → *θ*), the dynamics of coupled CANNs is written as,

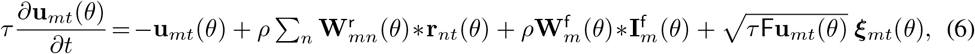

where **u**_*mt*_(*θ*) and **r**_*mt*_(*θ*) represent, respectively, the synaptic inputs and firing rates of neurons at time *t* in network *m* whose preferred value of stimulus feature *s*_*m*_ is *θ*. *τ* is the time constant, and *ρ* = *N*/*w*_*s*_ is the neuronal density covering the stimulus feature space. F is the Fano factor of the internal Poisson-like variability. 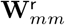 denotes the recurrent connection kernel between neurons in the same network, 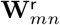 for *m* ≠ *n* is the reciprocal connection kernel between neurons from network *n* to network *m*, and 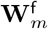 is the feedforward connection kernel. These kernels have the form of Gaussian function and are written as,

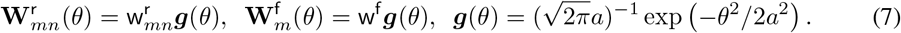

The symbol * denotes the convolution, i.e., **W**(*θ*) * **r**(*θ*) = ∫**W**(*θ* − *θ*′)**r**(*θ*′)*dθ*′, which implies the translation-invariant property of the connection pattern between neurons over the feature space, a key characteristic of CANNs. For simplicity, we assume the feedforward connection weight w^f^ is the same in different networks.

The relationship between the synaptic input and firing rate of neurons is modelled as divisive normalization [32, 33], which is given by

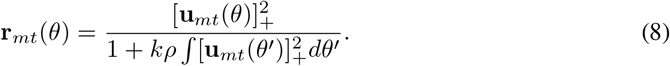

Here *k* determines the global inhibition strength and [·]_+_ is negative rectification. Divisive normalization is a canonical operation widely observed in the cortex and could be implemented via parvalbumin (PV) inhibitory neurons [34].

#### Feedforward input encoding the stimulus likelihood

In our model each network *m* receives one feedforward input 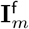, which conveys information about the latent stimulus feature *s*_*m*_ (Fig. 2A). Given the stimulus feature *s*_*m*_, 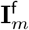 is modeled as independent Poisson spikes with Gaussian tuning (mean firing rate) (Fig. 2C). Mathematically, the probability of observing a particular value of 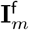 given *s*_*m*_ is,

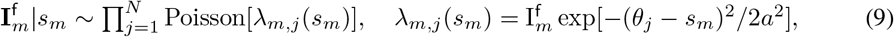

where *λ*_*m,j*_(*s*_*m*_) is the mean firing rate of the input component 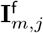 received by *j*-th neuron in network *m*, with 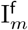 and *a* characterizing the peak input rate and tuning width, respectively. The above feedforward input has been widely used in previous neural coding studies, and satisfies the linear probabilistic population code proposed in [12]. Based on the Gaussian tuning and Poisson variability (Eq. 9), the likelihood of *s*_*m*_ given an observed 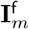 is also a Gaussian distribution (Fig. 2E), i.e., 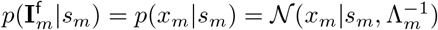, whose mean and precision are linear over 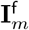 and are calculated to be

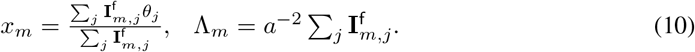

**Figure 2:**
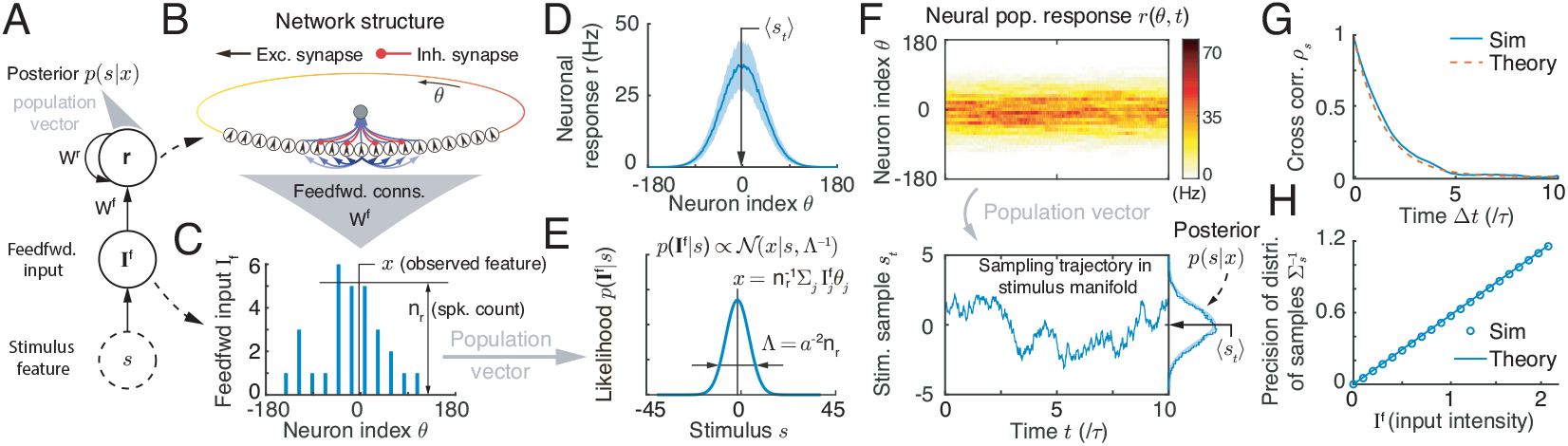
Langevin sampling of one dimensional stimulus feature in a CANN. (A) The information flow of the generative model. (B) The structure of a CANN. (C) The feedforward input is modeled as continuous approximation of Poisson spikes with Gaussian tuning over the stimulus feature. (D) The time averaged neural population responses, with the shaded region denoting standard deviation. (E) The likelihood is encoded by the feedforward input parametrically. (F) Neural population responses of the CANN over time (top); stimulus feature values sampled by the network dynamics (bottom left), whose distribution gives the posterior (bottom right; empirical: solid line, theory: shaded line). The theoretical value of the posterior is obtained from the feedforward input **I**^f^ by using Eq. (10) (see details in SI. Sec. 4.3). (G) Temporal correlation of sampling over time. (H) The precision (inverse of variance) of samples in equilibrium with the input intensity. Parameters can be found in SI. Sec. 4.

Comparing to Eq. 1, we see that the feedforward input 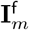 conveys the likelihood of the latent stimulus feature *s*_*m*_ parametrically.

#### Internal Poisson-like variability for reliable sampling

Implementing Langevin sampling in a network requires the network to generate additional internal variability not inherited from the feedforward inputs, which produces random walk on the log-posterior surface of the stimulus features (Eq. 4). Moreover, this internal variability should be Gaussian distributed with a white spectrum, so that the variance of samples (Eq. 5) matches the variance of the posterior (**Ω**^−1^ in Eq. 3).

Encoding the variance of the posterior is necessary for the brain to perceive the uncertainty of inputs [12, 17, 35]. To achieve Langevin sampling in our model, we consider that each network generates independent internal Poisson-like variability at the single neuron level (the last term in Eq. 6, e.g., [36–38]), which makes neural responses **u**_*m*_ and **r**_*m*_ exhibit the Poisson-like variability. As will be shown later, this Poisson-like variability in neural activities contributes to the required Gaussian-white variability in the stimulus features space (embedded in neural population responses), ensuring the realization of Langevin sampling. The Poisson-like variability provides a reliable source of variability to conduct sampling, as the Fano factor of neuronal responses only changes mildly across stimulus conditions [7, 12, 39, 40]. In reality, this internal Poisson-like (spiking) variability can naturally arise from the chaotic state of cortical circuit dynamics [40–44].

#### Network responses and decoding

Given the feedforward input **I**^f^, our theoretical analyses and simulations reveal that in the equilibrium, the mean neuronal responses of each network *m* have a Gaussian shape (Fig. 2D, i.e., [45, 46]),

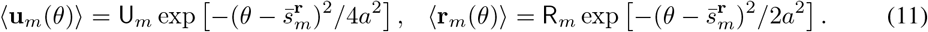

Here, U_*m*_ and R_*m*_ denotes the peak values of synaptic inputs and firing rate respectively. 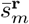 is the position of the Gaussian center on the stimulus feature space (Fig. 2D), which is the mean of stimulus feature samples in the network dynamics (see below). Because of the Gaussian tuning (Eq. 11) and the Poisson-like variability, given an instantaneous neuronal response **r**_*mt*_(*θ*) in network *m*, an instantaneous value of the stimulus feature can be efficiently read out using population vector [12, 47], i.e.,

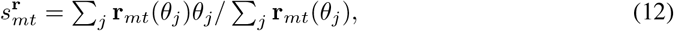

which is regarded as a sample of stimulus *s*_*m*_ (Eq. 14). Notably, the stimulus feature 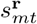 is read out based only on local neuronal responses in network *m*. Moreover, due to the divisive normalization operation in the network dynamics (Eq. 8), the Gaussian profile of neural responses (Eq. 11) can be well maintained even if the input disparity |*x*_*m*_ − *x*_*n*_| is large (Eq. 9), which ensures that stimulus features can always be efficiently read out by population vector.

### 3.2 Langevin sampling of stimulus features in coupled networks

We now elucidate how coupled CANNs can realize Langevin sampling of the posterior of high-dimensional stimulus features in a distributed manner. We apply perturbative analysis to derive the dynamics of stimulus features embedded in neural population responses (see details in SI. Sec. 3.2). For each network *m*, we consider that its instantaneous neural response is perturbed from its equilibrium mean, i.e., **u**_*mt*_(*θ*) = ⟨**u**_*m*_(*θ*)⟩ + *δ***u**_*mt*_(*θ*). The (unnormalized) eigenfunction of the perturbation *δ***u**_*mt*_(*θ*) corresponding to the change of stimulus feature *s*_*m*_ is derived as,

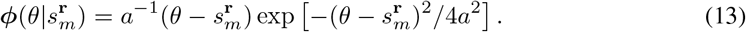

Previous studies have shown this eigenfunction has the largest eigenvalue for the perturbation of CANN state [45]. We project the dynamics of each network *m* onto the corresponding eigenfunction (Eq. 13), where the projection is simply the inner product between the eigenfunction and the network response, i.e., 〈***ϕ***, **u**_*mt*_〉 = ∫***ϕ***(*θ*)**u**_*mt*_(*θ*)*dθ*. After projection, we obtain the dynamics on the manifold of stimulus features embedded in neural responses (see details in SI 3.2),

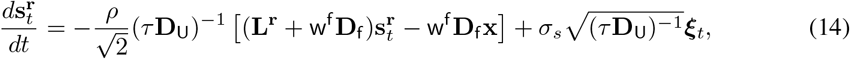

where 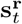 denotes the instantaneous positions of neural activity bumps at time *t*, which is regarded as a sample of stimulus features by the coupled networks (Eq. 12). **x** is the observed stimulus feature (Eq. 1) conveyed by the feedforward inputs (Eq. 9, Fig. 2E). **L^r^** is a generalized Laplacian matrix with 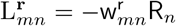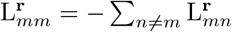, where R_*n*_ is the peak firing rate of network *n* (Eq. 11). w^f^ is a scalar variable denoting the feedforward connection strength (Eq. 7). **D**_U_ = diag(U_1_, U_2_, ···, U_*M*_) and 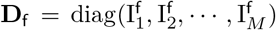 are diagonal matrices, denoting the peak value of the mean synaptic input, and the mean feedforward input in each network respectively (Eqs. 9 and 11). **D**_U_ can be analytically solved in our model (Eqs. S13-S14). Notably, after projection, the internal Poisson-like variability of neuronal activities (Eq. 6) becomes the Gaussian-white variability of stimulus features (the last term in Eq. 14), as required by Langevin sampling. The strength of Gaussian-white variability in the stimulus feature manifold is 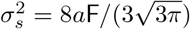, which is unchanged with respect to the feedforward input and network responses.

It is interesting to compare the dynamics on stimulus features (Eq. 14) with that of Langevin sampling (Eq. 4): both equations contain a Laplacian matrix (**L** or **L^r^**) representing the stimulus prior, a diagonal matrix (**Λ** or **D**_f_) representing the precision matrix of the likelihood, and Gaussian-white noises for producing a random walk. Thus, the dynamics of coupled CANNs in effect realizes Langvein sampling in the manifold of stimulus features. The equilibrium distribution of stimulus features **s^r^** in Eq. (14) is a multivariate Gaussian, denoted as 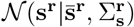, whose mean 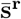 and covariance of 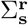 satisfy two conditions (for details, see SI. Sec. 2),

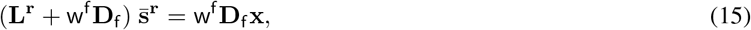

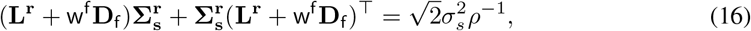

where 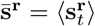, and 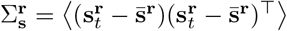. By setting the equilibrium mean and covariance of stimulus features sampled by the network to be equal to that of the posterior of latent stimulus features, i.e., 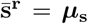 and 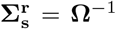 (see Eq. 3), we get the required network connections. It can be checked that 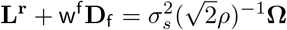 can make 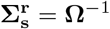 hold. Substituting this into Eqs. (15–16), we get the network connections for realizing Langevin sampling of the posterior of stimulus features, which are,

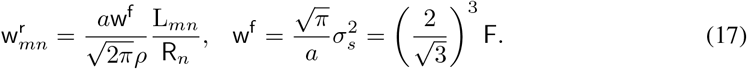

Interestingly, we see that the reciprocal connections 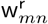 and 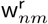 between networks *n* and *m* encode the prior (the correlation) L_*mn*_ between stimulus features *s*_*m*_ and *s*_*n*_ (Eq. 1). Moreover, although the prior precision matrix is symmetric, i.e., L_*mn*_ = L_*nm*_, the couplings 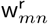 and 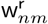 can be asymmetric, which is biologically more plausible. When L_*mn*_ = 0 for *m* ≠ *n*, the prior *p*(**s**) degenerates into uniform, and the reciprocal connections 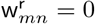, for *m* ≠ *n*. In such a case, each network individually samples its marginal posterior which is proportional to the likelihood.

In summary, we have shown that the model of coupled CANNs with appropriate feedforward inputs (conveying the likelihoods, Eq. 9), appropriate reciprocal connections (encoding the prior, Eq. 17), and appropriate internal variability (producing random walks, Eq. 14) can implement Langevin sampling on the manifold of high-dimensional latent stimulus features. Further, this is done in a distributed manner, in term of that each network infers the marginal posterior of one stimulus feature. Finally, the sampled stimulus features can be read out easily by using a linear decoder (population vector) and separately from local neural activities at each network (see Eq. 12).

## 4 Simulation experiments

We carry out simulations to validate the above theoretical analysis. We first consider a single CANN (equivalent to an uncoupled CANN; Fig. 2A-B). In this case each network *m* independently infers a posterior *p*(*s*_*m*_|*x*_*m*_) based on the likelihood *p*(*x*_*m*_|*s*_*m*_) and the uniform stimulus prior. We present a constant feedforward input 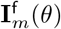 generated by Eq. 9 to the network, and then evaluate the sampling performance of the network (more details can be found in SI. Sec. 4). Firstly, we observe that due to the internal variability, the neural population responses fluctuate over time (Fig. 2F), whose time-averaged profile has a Gaussian shape and individual neuronal responses exhibit Poisson-like variability (Eq. 11, Fig. 2D). This guarantees that the instant sample of stimulus feature 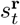 can be efficiently read out from the instantaneous neuronal response **r**_*t*_ using population vector (Eq. 12). Secondly, we observe that the network performs Langevin sampling. As shown in Fig. 2F (bottom), the stimulus feature 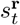 sampled by the network shows random walk behavior over time, a characteristic of Langevin sampling. Moreover, the temporal correlation of sampling agrees with the theoretical prediction of Langevin sampling, i.e., 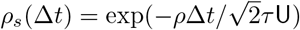 (Fig. 2G). Thirdly, we observe that by setting the feedforward weight as in Eq. 17, the equilibrium distribution of sampled stimulus feature (i.e., the distribution of sequential samples in equilibrium, Fig. 2F, bottom) is consistent with the posterior (Fig. 2H). Thus, a single CANN with internal Poisson-like variability can implement Langevin sampling on the stimulus feature manifold.

Next, we demonstrate that coupled CANNs implement Langevin sampling of the posterior of high-dimensional stimulus features. We first consider a model of two coupled networks (Fig. 3A), in which the stimulus manifold is a torus, with each circle representing a stimulus feature (Fig. 3B). Similar to the case of a single CANN, neuronal responses at each network exhibits Langevin sampling behaviors due to the internal variability. The instant stimulus estimate 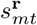 can be read out using population vector based only on the local instantaneous neural activity **r**_*mt*_ in network *m* (Eq. 12, Fig. 3A). This implies that a neural decoder only needs to access neural activity locally rather than over distributed brain areas. We find that the network dynamics is performing Langevin sampling on the manifold of torus (Fig. 3C), and that the equilibrium distribution of samples agrees with the theoretical prediction (Fig. 3D, see details in SI. Sec. 4.3). In particular, the distribution of sampled feature values at each individual network agrees with the marginal posterior of the corresponding stimulus feature, demonstrating that our model is performing inference in a distributed manner. Furthermore, we confirm that the reciprocal coupling between two networks encodes the prior of stimulus features (i.e., the precision matrix, Fig. 3E, Eq. 17).

**Figure 3:**
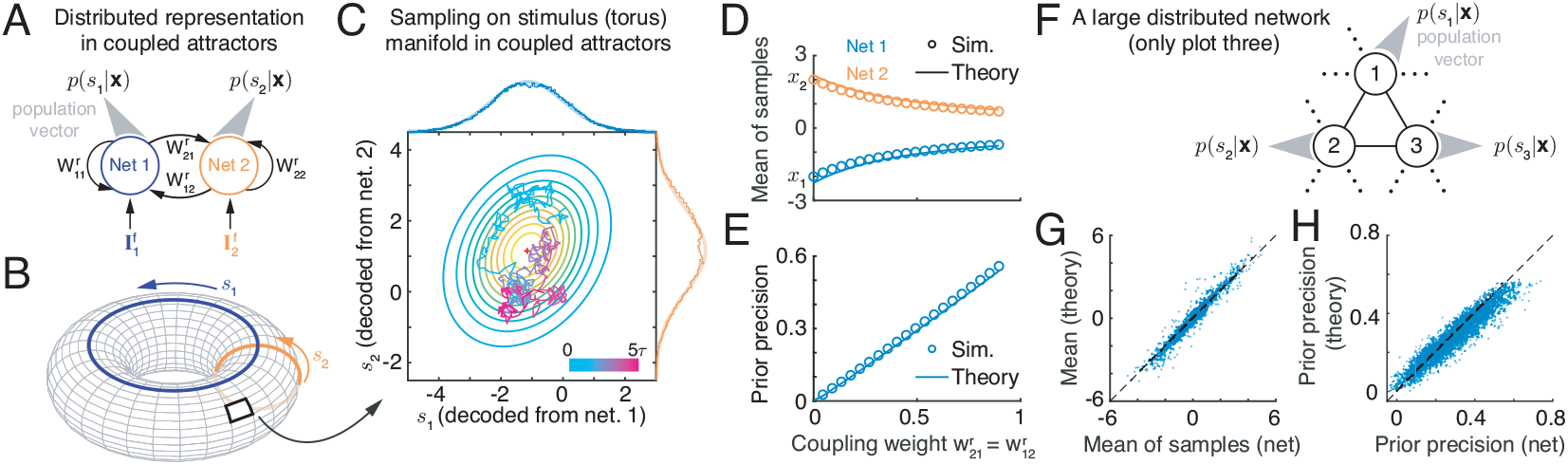
Distributed Langevin sampling in coupled CANNs. (A) The structure of two-coupled CANNs. Each network receives a feedforward input conveying the likelihood of stimulus feature and meanwhile is reciprocally connected to other networks. (B) The torus manifold of two-dimensional stimulus features embedded in two coupled CANNs. (C) The model of two-coupled CANNs implements Langevin sampling of the posterior of stimulus features, for each network inferring the corresponding marginal posterior. Solid ellipses: the posterior by theory. Colors of the trajectory indicate the time elapsed. Marginal plot: marginal posterior (empirical: solid line; theory: shaded line). (D) The mean of samples in each network is consistent with the mean of the posterior. (E) The coupling strengths encode the precision of the prior (for detailed calculations, see SI. Sec. 4.4). (F) A large network contains up to ten coupled CANNs. (G-H) The comparisons of the mean of sampling (G) and the prior precision stored in the network (H) with theoretical predictions. Each point is a result obtained under a combination of different inputs, different connection weights, and different realizations of the network numbers. Parameters can be found in SI. Sec. 4.

Our model is robust and scalable to arbitrary number of networks for processing arbitrary dimension of stimulus features. To demonstrate this, we simulate a model containing up to ten coupled CANNs (Fig. 3F), and verify that indeed each network is able to implement Langevin sampling of the marginal posterior of the corresponding stimulus feature (Fig. 3G-H).

## 5 Conclusions and Discussions

In this study, we propose that coupled CANNs with appropriate structures are able to implement distributed, sampling-based inference to approximate the multivariate posterior of high-dimensional stimulus features. We elucidate that the dynamics of coupled CANNs in effect realizes Langevin sampling in the manifold of stimulus features, where the feedforward inputs convey the likelihood, the reciprocal connections encode the priors of stimulus association, and the internal Poisson variability of the neurons produce random walks for sampling. Our model achieves sampling-based inference in a distributed manner, in term of that each network infers the marginal posterior of the corresponding stimulus feature. Furthermore, the sampled stimulus features can be read out easily using population vector based only on local neural activities at each network.

Our model is different from others in the literature for implementing Bayesian inference. Compared to PPC and its extended version to coupling networks [12, 13], our model is distinguished in that: 1) the inference and representation of the posterior in our model is sampling-based, rather than parametrically represented; 2) our model is stochastic, which generates internal Poisson variability (not as inherited from the noisy feedforward inputs as in PPC) to trigger random walks of sampling stimulus features; 3) the marginalization via sampling in our model collectively emerges from the stochastic dynamics of coupled networks, rather than through message-passing mediated by nonlinear couplings between networks [13]. Compared to other sampling-based inference models (e.g., [14–20]), our model considers that sampling occurs on the manifold of stimulus features embedded in neuronal population responses, where stimulus features are the variables of interest. Furthermore, we develop a concrete neural circuit model to implement this inference, and demonstrate that the Poisson (spiking) variability of the neurons contributes to the Gaussian-white variability of stimulus features necessary for Langevin sampling. A previous study also employed coupled CANNs to realize distributed multisensory integration [46], but their way of realizing Bayesian inference is not sampling-based, but rather being a statistical result by averaging over input trials. Overall, we have proposed a novel model which integrates the strengths of PPC and sampling-based codes in a unified framework to achieve high-dimensional Bayesian inference efficiently.

Nevertheless, there are some detailed issues needing to be addressed in the future. Langevin sampling is known to be slow especially in a high-dimensional space [48]. In our model, sampling occurs on the manifold of stimulus features, whose dimension (equalling to the number of coupled networks) is already smaller by orders of magnitude compared to the dimension of neuronal responses (equalling to the number of neurons), but still the sampling speed may be not quick enough in practice. Our analysis shows that the temporal correlation of sampling in our model is 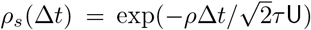 (Fig. 2G), where the decaying time constant 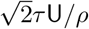 (whose inverse quantifying the sampling speed) increases with the neural activity U (Eq. 11). To further speed up sampling, a potential strategy is to include short-term synaptic plasticity [49], which increases the transition probability between network states [50] and has the potential to generate effective anti-symmetric connections between networks to speed up sampling as proposed in [16]. We will study this issue in future work.

## Acknowledgments

This work is supported by National Science Foundation (1816568), the National Institutes of Health (Grants 1U19NS107613-01 and R01EB026953), the Vannevar Bush Faculty (Fellowship N00014-18-1-2002), and the Simons Foundation Collaboration on the Global Brain.

