## Supplementary information for "Distributed Sampling-based Bayesian Inference in Coupled Neural Circuits"

#### 1 The derivation of Lyapunov equation

We present the mathematical derivation of Lyapunov equation whose solution denotes the equilibrium covariance of a Langevin dynamics (Ornstein–Uhlenbeck process). Given a general Langevin dynamics,

$$d\mathbf{s}(t) = \mathbf{M}[\mathbf{x} - \mathbf{s}(t)] + \boldsymbol{\beta} d\mathbf{W}_t, \tag{S1}$$

where $\mathbf{x}$ and $\mathbf{s}$ are both $N$-by-1 vectors, and $\mathbf{W}_t$ denotes a $N$-dim standard Wiener process with the elements independent with each other, i.e., $\langle d\mathbf{W}_t(i) d\mathbf{W}_{t'}(j)\rangle = \delta_{ij}\delta(t-t')$, its solution of $\mathbf{s}(t)$ can be calculated as,

$$\mathbf{s}(t) = e^{-\mathbf{M}t}\left[\mathbf{s}(0) + \int_0^t e^{\mathbf{M}t'}\mathbf{x}dt' + \int_0^t e^{\mathbf{M}t'}\boldsymbol{\beta} d\mathbf{W}_{t'}\right]. \tag{S2}$$

Then the covariance $\boldsymbol{\Sigma}_\mathbf{s}(t) = \langle (\mathbf{s}(t) - \langle \mathbf{s}(t)\rangle)(\mathbf{s}(t) - \langle \mathbf{s}(t)\rangle)^\top \rangle$ averaged across trials at time $t$ can be derived as

$$\boldsymbol{\Sigma}_\mathbf{s}(t) = e^{-\mathbf{M}t}\left[\boldsymbol{\Sigma}_\mathbf{s}(0) + \left\langle \left(\int_0^t e^{\mathbf{M}t'}\boldsymbol{\beta} d\mathbf{W}_{t'}\right)\left(\int_0^t e^{\mathbf{M}t'}\boldsymbol{\beta} d\mathbf{W}_{t'}\right)^\top \right\rangle\right]\left(e^{-\mathbf{M}t}\right)^\top. \tag{S3}$$

Denote by $\mathbf{A}(t)$ the terms inside the bracket in above equation, the evolution of covariance $\Sigma_\mathbf{s}(t)$ can be converted into a differential equation by taking the derivative over time $t$,

$$\begin{aligned}
\frac{d\boldsymbol{\Sigma}_\mathbf{s}(t)}{dt} &= \frac{d}{dt}\left[e^{-\mathbf{M}t}\mathbf{A}(t)\left(e^{-\mathbf{M}t}\right)^\top\right], \\
&= \frac{de^{-\mathbf{M}t}}{dt}\mathbf{A}(t)\left(e^{-\mathbf{M}t}\right)^\top + e^{-\mathbf{M}t}\frac{d\mathbf{A}(t)}{dt}\left(e^{-\mathbf{M}t}\right)^\top + e^{-\mathbf{M}t}\mathbf{A}(t)\frac{d\left(e^{-\mathbf{M}t}\right)^\top}{dt}, \\
&= -\mathbf{M}e^{-\mathbf{M}t}\mathbf{A}(t)\left(e^{-\mathbf{M}t}\right)^\top + \boldsymbol{\beta}\boldsymbol{\beta}^\top - e^{-\mathbf{M}t}\mathbf{A}(t)\left(e^{-\mathbf{M}t}\right)^\top\mathbf{M}^\top, \\
&= -\mathbf{M}\Sigma_\mathbf{s}(t) + \boldsymbol{\beta}\boldsymbol{\beta}^\top - \boldsymbol{\Sigma}_\mathbf{s}(t)\mathbf{M}^\top.
\end{aligned} \tag{S4}$$

Hence the covariance $\Sigma_\mathbf{s}(t)$ in the equilibrium state $t \to \infty$ is

$$\mathbf{M}\boldsymbol{\Sigma}_\mathbf{s} + \boldsymbol{\Sigma}_\mathbf{s}\mathbf{M}^\top = \boldsymbol{\beta}\boldsymbol{\beta}^\mathrm{T}. \tag{S5}$$

This is the Lyapunov equation shown in Eq. (16) in main text, which characterizes the equilibrium covariance of the Langevin dynamics.

Preprint. Under review.

#### 2 The equilibrium distribution of Langevin sampling

We provides the math details of calculating the equilibrium distribution of Langevin sampling as shown in Eq. (4).

$$\frac{d\mathbf{s}_t}{dt} = -(2\tau_L)^{-1}\left[(\mathbf{L}+\mathbf{\Lambda})\mathbf{s}_t - \mathbf{\Lambda}\mathbf{x}\right] + \sqrt{\tau_L^{-1}}\boldsymbol{\xi}_t, \tag{S6}$$

Since the noises $\boldsymbol{\xi}_t$ are Gaussian white with zero mean, the mean of samples $\bar{\mathbf{s}}_t = \langle \mathbf{s}_t \rangle$ in equilibrium could be obtained right after taking the average of above equation and setting the derivative term into zero,

$$\bar{\mathbf{s}}_t = (\mathbf{L}+\mathbf{\Lambda})^{-1}\mathbf{\Lambda}\mathbf{x} = \mathbf{\Omega}^{-1}\mathbf{\Lambda}\mathbf{x}. \tag{S7}$$

Meanwhile, the equilibrium covariance could be obtained by solving the Lyapunov equation (Eq. S5). Comparing the Eqs. (S6 and S1), the Lyapunov equation governing the equilibrium covariance in Eq. (S6) is,

$$\left[-(2\tau_L)^{-1}\mathbf{\Omega}\right]\mathbf{\Sigma_s} + \mathbf{\Sigma_s}\left[-(2\tau_L)^{-1}\mathbf{\Omega}\right]^\top = -\tau_L^{-1}\mathbf{I}. \tag{S8}$$

It can be checked that $\mathbf{\Sigma_s} = \mathbf{\Omega}^{-1}$ is the solution of above Lyapunov equation, suggesting the Langevin sampling indeed approximates the posterior.

In addition, the network dynamics on the stimulus feature manifold (Eq. 14) is also a Langevin dynamics, whose equilibrium mean and covariance can be solved similarly as shown in Eqs. (S6-S8), which yields the Eqs. (15 and 16) in main text.

#### 3 Theoretical analysis the attractor network

##### 3.1 Mean neuronal responses of each attractor network

We firstly analyze the mean population responses in the equilibrium in the coupled attractor networks, i.e., the dynamics without the Poisson-like variability in Eq. (11). Because the recurrent connection kernel $\mathbf{W}_{mn}(\theta)$ is a gaussian function, and the convolution of two Gaussian functions are still Gaussian, we propose the ansatz that the equilibrium mean population response of each attractor network in coupled networks has Gaussian profile approximately, as Eq. (11) in main text,

$$\langle \mathbf{u}_m(\theta)\rangle \approx \mathsf{U}_m \exp[-(\theta-\bar{s}^{\mathbf{r}}_m)^2/4a^2], \quad \langle \mathbf{r}_m(\theta)\rangle \approx \mathsf{R}_m \exp[-(\theta-\bar{s}^{\mathbf{r}}_m)^2/2a^2], \tag{S9}$$

Substituting Eq. (S9) into the network dynamics (Eq. 6), the recurrent inputs in the network can be computed as

$$\begin{aligned}\rho\mathbf{W}^{\mathsf{r}}_{mn}(\theta) * \mathbf{r}_{nt}(\theta) &= \frac{\rho \mathsf{w}^{\mathsf{r}}_{mn}\mathsf{R}_n}{\sqrt{2\pi}a}\int e^{-(\theta-\theta')^2/2a^2-(\theta'-\bar{s}^{\mathbf{r}}_n)^2/2a^2}d\theta', \\ &= \frac{\rho \mathsf{w}^{\mathsf{r}}_{mn}\mathsf{R}_n}{\sqrt{2}}e^{-(\theta-\bar{s}^{\mathbf{r}}_n)^2/4a^2}.\end{aligned} \tag{S10}$$

And similarly the feedforward input received by network $m$ is,

$$\rho\mathbf{W}^{\mathsf{f}}_m(\theta) * \mathbf{I}^{\mathsf{f}}_m(\theta) = \frac{\rho \mathsf{w}^{\mathsf{f}}_m \mathrm{I}^{\mathsf{f}}_m}{\sqrt{2}}e^{-(\theta-x_m)^2/4a^2}. \tag{S11}$$

Hence the equilibrium mean response given the Gaussian ansatz (Eq. S9) is,

$$\mathsf{U}_m e^{-(\theta-\bar{s}^{\mathbf{r}}_m)^2/4a^2} \approx \frac{\rho}{\sqrt{2}}\Big[\sum_n \mathsf{w}^{\mathsf{r}}_{mn}\mathsf{R}_n e^{-(\theta-\bar{s}^{\mathbf{r}}_n)^2/4a^2} + \mathsf{w}^{\mathsf{f}}_m \mathrm{I}^{\mathsf{f}}_m e^{-(\theta-x_m)^2/4a^2}\Big]. \tag{S12}$$

From Eq. (S12), if the feature of every feedforward input $x_m$ is exactly the same, i.e., $x_m \equiv x$, or without any feedforward input ($\mathrm{I}_m = 0$), the location of network responses will be exactly the same, i.e., $s^{\mathbf{r}}_m \equiv s^{\mathbf{r}}$, and then the mean response $\langle \mathbf{u}_m(\theta)\rangle$ will be a perfect Gaussian. In this case, the height of the population resposne can be solved exactly as,

$$\mathsf{U}_m = \frac{\rho}{\sqrt{2}}\Big(\sum_n \mathsf{w}^{\mathsf{r}}_{mn}\mathsf{R}_n + \mathsf{w}^{\mathsf{f}}_m \mathrm{I}^{\mathsf{f}}_m\Big), \tag{S13}$$

$$\mathsf{R}_m = \frac{\mathsf{U}_m^2}{1+\sqrt{2\pi}k\rho a \mathsf{U}_m^2}, \tag{S14}$$

where Eq. (S14) is obtained through submitting the Gaussian ansatz into the the divisive normalization (Eq. 8).

When the input disparity $|x_m - x_n|$ is large, the mean response will be distorted from a perfect Gaussian profile. However, due to the translation-invariant recurrent connection (convolution in stimulus feature space) and the nonlinear divisive normalization, the mean response of each network is still a bell-shape profile and never exhibits multimodel as confirmed by our simulation. And then the Gaussian ansatz still works well enough.

##### 3.2 The network dynamics on the stimulus manifold

We are interested in analyzing the network dynamics on the stimulus manifold. Using perturbative analysis, we consider the actual mean response is the perturbed version from the perfect Gaussian profile in equilibrium state ($\langle \mathbf{u}_m(\theta)\rangle$ in Eq. S9). Previous theoretical study indicates the (unnormalized) eigenfunction corresponding to the perturbation (change) on the stimulus $s_m$'s manifold is proportional to the derivative of the mean response $\langle \mathbf{u}_m(\theta)\rangle$ over the stimulus $s_m$ [1],

$$\boldsymbol{\phi}(\theta|\bar{s}^{\mathbf{r}}_m) \propto \frac{d}{d\bar{s}^{\mathbf{r}}_m}\langle \mathbf{u}_m(\theta)|\bar{s}^{\mathbf{r}}_m\rangle = a^{-1}(\theta - \bar{s}^{\mathbf{r}}_m)e^{-(\theta-\bar{s}^{\mathbf{r}}_m)^2/4a^2}. \tag{S15}$$

And then we could project the high dimensional network dynamics (Eq. 6) onto above eigenfunction (Eq. S15) to get the dynamics on the stimulus manifold. Projecting a function $f(\theta)$ onto the normalized eigenfunction $\boldsymbol{\phi}(\theta|s^{\mathbf{r}}_m)$ corresponds to compute the inner product between the two and normalizing by the norm of the eigenfunction,

$$\frac{\langle f(\theta), \boldsymbol{\phi}(\theta|\bar{s}^{\mathbf{r}}_m)\rangle}{\langle \boldsymbol{\phi}(\theta|\bar{s}^{\mathbf{r}}_m), \boldsymbol{\phi}(\theta|\bar{s}^{\mathbf{r}}_m)\rangle} = \frac{\int f(\theta)\boldsymbol{\phi}(\theta|\bar{s}^{\mathbf{r}}_m)d\theta}{\int \boldsymbol{\phi}(\theta|\bar{s}^{\mathbf{r}}_m)^2 d\theta} = \frac{1}{\sqrt{2\pi}a}\int f(\theta)\boldsymbol{\phi}(\theta|\bar{s}^{\mathbf{r}}_m)d\theta. \tag{S16}$$

Let's show some detailed math derivation of the projection of the dynamics of network $m$ onto eigenfunction $\boldsymbol{\phi}(\theta|s^{\mathbf{r}}_m)$. Substituting the Gaussian ansatz (Eq, S9) into network dynamics (Eq. 6),

$$\begin{aligned}
&\tau\frac{\mathsf{U}_m}{2a}\frac{ds^{\mathbf{r}}_{mt}}{dt}\frac{\theta - s^{\mathbf{r}}_{mt}}{a}e^{-(\theta-s^{\mathbf{r}}_{mt})^2/4a^2},\\
&= -\mathsf{U}_m e^{-(\theta-s^{\mathbf{r}}_{mt})^2/4a^2} + \frac{\rho}{\sqrt{2}}\sum_n \mathsf{w}^{\mathsf{r}}_{mn}\mathsf{R}_n e^{-(\theta-s^{\mathbf{r}}_{nt})^2/4a^2},\\
&\quad + \frac{\rho}{\sqrt{2}}\mathsf{w}^{\mathsf{f}}_m \mathrm{I}^{\mathsf{f}}_m e^{-(\theta-x_m)^2/4a^2} + \sqrt{\tau \mathsf{F}\mathsf{U}_m}e^{-(\theta-s^{\mathbf{r}}_{mt})^2/8a^2}\boldsymbol{\xi}_{mt}.
\end{aligned} \tag{S17}$$

Projecting above dynamics onto the eigenfunction $\boldsymbol{\phi}(\theta|s^{\mathbf{r}}_m)$,

$$\begin{aligned}
\tau\frac{\mathsf{U}_m}{2a}\frac{ds^{\mathbf{r}}_{mt}}{dt} &= \frac{\rho}{2\sqrt{2}a}\sum_n \mathsf{w}^{\mathsf{r}}_{mn}\mathsf{R}_n(s^{\mathbf{r}}_{nt} - s^{\mathbf{r}}_{mt})e^{-(s^{\mathbf{r}}_n - s^{\mathbf{r}}_m)^2/8a^2},\\
&\quad + \frac{\rho}{2\sqrt{2}a}\mathsf{w}^{\mathsf{f}}_m \mathrm{I}^{\mathsf{f}}_m(x_m - s^{\mathbf{r}}_{mt})e^{-(x_m - s^{\mathbf{r}}_{mt})^2/8a^2} + \left(\frac{2\mathsf{F}\tau\mathsf{U}_m}{3\sqrt{3\pi}a}\right)^{1/2}\xi_{mt}.
\end{aligned} \tag{S18}$$

Suppose the difference of the position of networks is small enough compared with the tuning width $a$, i.e., $|s^{\mathbf{r}}_n - s^{\mathbf{r}}_m| \ll 4a$, the exponential terms in above equation can be ignored for simplicity. Reorganize the terms in above equation,

$$\begin{aligned}
\frac{ds^{\mathbf{r}}_{mt}}{dt} &= \frac{\rho}{\sqrt{2}\tau\mathsf{U}_m}\Big[\sum_n \mathsf{w}^{\mathsf{r}}_{mn}\mathsf{R}_n(s^{\mathbf{r}}_{nt} - s^{\mathbf{r}}_{mt}) + \mathsf{w}^{\mathsf{f}}_m\mathrm{I}^{\mathsf{f}}_m(x_m - s^{\mathbf{r}}_{mt})\Big],\\
&\quad + \left(\frac{8a\mathsf{F}}{3\sqrt{3\pi}}\right)^{1/2}(\tau\mathsf{U}_m)^{-1/2}\xi_{mt}.
\end{aligned} \tag{S19}$$

Denoting $\sigma_s^2 = 8a\mathsf{F}/(3\sqrt{3\pi})$, and writing the above dynamics in matrix form, we get the Eq. (14) in the main text.

### 4 Details of network simulation

#### 4.1 Network parameter and simulation details

The width each stimulus feature space is $\mathsf{w}_s = 360°$ for simplicity. And each attractor network $m$ contains $N = 180$ neurons which are uniformly distributed in the space of $s_m$. And hence the neuronal density $\rho = N/\mathsf{w}_s = 0.5$. The connection width $a = 40°$, which will make the width of population response and the tuning of single neurons the same as $40°$, consistent with V1 data. The synaptic time constant $\tau$ is rescaled to 1 as a dimensionless number. The Fano factor $\mathsf{F}$ of synaptic input $\mathbf{u}_m(\theta)$ is set to $0.5$, which will make the average Fano factor of firing rate $\mathbf{r}_m(\theta)$ around 1 (Fig. 2D). And the global connection strength $k = 5 \times 10^{-4}$ for each attractor network (Eq. 8), which makes the peak firing rate $r_m(\theta)$ of each network saturate at the level around 50Hz, consistent with typical experimental observations.

To scale the connection strength in the network model, we find the smallest recurrent connection strength $\mathsf{w}_c$ under which a single attractor network could maintain a persistent activity after switching off the feedforward input. Cutting off all couplings between networks (setting $\mathsf{w}^{\mathsf{r}}_{mn} = 0$ for $m \neq n$), and removing feedforward input (setting $\mathrm{I}^{\mathsf{f}}_m = 0$), $\mathsf{w}^{\mathsf{r}}_c$ can be solved by combining Eqs. (S13 and S14),

$$\mathsf{w}^{\mathsf{r}}_c = 2\sqrt{2}(2\pi)^{1/4}\sqrt{ka\rho} \approx 0.896. \tag{S20}$$

And then the recurrent strength within an attractor network $\mathsf{w}^{\mathsf{r}}_{mm}$, and the coupling stregnth across networks $\mathsf{w}^{\mathsf{r}}_{mn}$ ($m \neq n$) will be both set as values relative to $\mathsf{w}^{\mathsf{r}}_c$.

Similarly, the peak rate of feedforward input, i.e., $\mathrm{I}^{\mathsf{f}}_m$, is also scaled with respect to $\mathsf{U}_c$

$$\mathsf{U}_c = \frac{\mathsf{w}^{\mathsf{r}}_c}{2\sqrt{\pi}ka} \approx 12.632, \tag{S21}$$

where $\mathsf{U}_c$ is the peak response of the synaptic input $\mathbf{u}_m(\theta)$ can be maintained by a single continuous attractor network when $\mathsf{w}^{\mathsf{r}} = \mathsf{w}^{\mathsf{r}}_c$ and the feedforward input is switched off.

For simplicity, the feedforward input $\mathbf{I}^{\mathsf{f}}_m$ in the simulation is set to be a smooth Gaussian function (corresponding to its mean averaged over trial) in simulation,

$$\mathbf{I}^{\mathsf{f}}_m = \mathrm{I}^{\mathsf{f}}_m \exp[-(\theta_j - x_m)^2/2a^2].$$

The network is simulated by using Euler method with time step $\Delta t = 0.01\tau$. To estimate the distribution of stimulus samples, we simulate the network model for a period of $600\tau$, and then the initial responses in the first $100\tau$ will be discarded and not included in estimating the equilibrium distribution of samples.

Following is the extra parameters used in Fig. 2 and 3.

| | |
|---|---|
| Fig. 2: | $\mathrm{I}^{\mathsf{f}}_m \in [0, 2\mathsf{U}_c]$ with a step of $0.1\mathsf{U}_c$, $\mathsf{w}^{\mathsf{r}} = 0.5\mathsf{w}^{\mathsf{r}}_c$, $x = 0°$. |
| Fig. 3D-E: | $\mathrm{I}^{\mathsf{f}}_m = \mathsf{U}_c$, $\mathsf{w}^{\mathsf{r}} = 0.5\mathsf{w}^{\mathsf{r}}_c$, $\mathsf{w}^{\mathsf{r}}_{12} = \mathsf{w}^{\mathsf{r}}_{21} \in [0, 2\mathsf{w}^{\mathsf{r}}]$ with a step of $0.5\mathsf{w}^{\mathsf{r}}$, and $x_1 = -2°$, $x_2 = 2°$. |
| Fig. 3G-H: | The network number $N \in [2, 10]$. $\mathrm{I}^{\mathsf{f}}_m \sim \mathcal{U}[0, 1.5\mathsf{U}_c]$, $\mathsf{w}^{\mathsf{r}}_{mn} \sim \mathcal{U}[0, \mathsf{w}^{\mathsf{r}}_c]$, $x_m \sim \mathcal{U}[-5, 5]$. For each network number, $\mathrm{I}^{\mathsf{f}}_m$, $\mathsf{w}^{\mathsf{r}}_{mn}$ and $x_m$ are randomly sampled for 50 times. Given each combination of network parameters, the network is simulated for $600\tau$. |

$\mathcal{U}[a, b]$ denotes a uniform distribution ranging from $a$ to $b$.

##### 4.2 Decoding the instantaneous stimulus samples

The instantaneous stimulus sample $s^{\mathbf{r}}_{mt}$ could be *locally* read out from the instantaneous population response $\mathbf{r}_{mt}$ at network $m$ by population vector (Eq. 10), i.e.,

$$s^{\mathbf{r}}_{mt} = \frac{\sum_j \mathbf{r}_{mt}(j)\theta_j}{\sum_j \mathbf{r}_{mt}(j)}. \tag{S22}$$

Note that in the network simulation, we could read out a trajectory of stimulus sample $s_{mt}$ from the spatiotemporal population response $\mathbf{r}_{mt}$, which is in constrat to only one stimulus estimate is read out from the neuronal response averaged across time in a trial in typical experiments. We collect the trajectory of stimulus samples $\mathbf{s}^{\mathbf{r}}_t$, with each individually read out from each network, and then compute the mean and covariance of these samples (corresponding to the numerical solution of Eqs. 15 and 16), i.e,

$$p(\mathbf{s}^{\mathbf{r}}) = \mathcal{N}(\mathbf{s}^{\mathbf{r}}|\bar{\mathbf{s}}^{\mathbf{r}}, \mathbf{\Sigma}_{\mathbf{s}}) \tag{S23}$$

which is supposed to approximate the posterior of the stimulus features, i.e. $p(\mathbf{s}|\mathbf{x})$ in Eq. (3).

##### 4.3 Evaluating the mean of samples in coupled networks

If the coupled networks indeed perform sampling-based inference, the mean of samples $p(\mathbf{s}^{\mathbf{r}})$ should satisfy the same property with the posterior mean (Eq. 3), i.e.,

$$\bar{\mathbf{s}}^{\mathbf{r}} = \mathbf{\Sigma}^{\mathbf{r}}_{\mathbf{s}} \mathbf{\Lambda} \mathbf{x}, \tag{S24}$$

where $\mathbf{\Sigma}_{\mathbf{s}}$ is the covariance of stimulus samples (Eq. S23). Hence Eq. (S24) can be used to verify whether the mean of samples correctly takes the uncertainty information ($\mathbf{\Sigma}_{\mathbf{s}}$).

In network simulation, the likelihood precision $\mathbf{\Lambda}$ can be directly decoded from feedforward inputs by using population vector (Eq. 10), because the feedforward input $\mathbf{I}^{\mathsf{f}}_m$ received by network $m$ averaged over trials satisfies the distribution specified in Eq. (9). For simplicity, the feedforward input $\mathbf{I}^{\mathsf{f}}_m$ is set to be a smooth Gaussian function (corresponding to its mean averaged over trial) in simulation,

$$\mathbf{I}^{\mathsf{f}}_m = \mathrm{I}^{\mathsf{f}}_m \exp[-(\theta_j - x_m)^2/2a^2].$$

And then the likelihood of $s_m$ given a snapshot of feedforward input $\mathbf{I}^{\mathsf{f}}_m$ can be read out to be,

$$p(s_m|\mathbf{I}^{\mathsf{f}}_m) = \mathcal{N}(s_m|x_m, \Lambda_m^{-1}), \tag{S25}$$

where the position of $\mathbf{I}_{\mathsf{f}m}$ specifies the mean of likelihood, $x_m$. And the precision of likelihood is (Eq. 10),

$$\begin{aligned}\Lambda_m &= a^{-2}\frac{1}{\Delta\theta}\sum_j \mathbf{I}^{\mathsf{f}}_m(\theta_j)\Delta\theta, \\ &\approx a^{-2}\rho \int \mathbf{I}^{\mathsf{f}}_m(\theta)d\theta = \sqrt{2\pi}\rho a^{-1}\mathrm{I}^{\mathsf{f}}_m,\end{aligned} \tag{S26}$$

where $\rho = 1/\Delta\theta = N/\mathrm{w}_s$ represents the density of neurons covering the stimulus feature space $s_m$. Eq. (S26) indicates the precision of the likelihood of $s_m$ is proportional to the peak firing rate of feedforward input $\mathrm{I}^{\mathsf{f}}_m$.

To verify the Eq. (S24), we read out the trajectory of stimulus samples from neuronal responses (Eq. S22), and then the distribution of stimulus samples can be estimated (Eq. S23). Meanwhile, the likelihood precision $\mathbf{\Lambda}$ can be decoded from feedforward inputs (Eq. S26). Then we substitute the decoded $\mathbf{\Sigma}^{\mathbf{r}}_{\mathbf{s}}$ and $\mathbf{\Lambda}$ into Eq. (S24) to get the prediction of the mean of sample, denoted by $\langle \mathbf{s}^{\mathbf{r}} \rangle_{\mathrm{pred}}$, and then it will be compared to the actual mean of samples $\langle \mathbf{s}^{\mathbf{r}} \rangle$. Fig. 3D presents the comparison and shows the mean of samples computed by network indeed correctly take the uncertainty information.

##### 4.4 Verifying the prior representation in coupled networks

Apart from verifying whether the mean of stimulus samples in network model consistent with the theoretical prediction (Eq. S24), we also verify whether the prior information is stored in the network as suggested by our theoretical derivation (Eq. 17).

To do this, given a connection weight matrix ($\mathbf{W}$ in Eq. 7), the network model is simulated and the distribution of samples $p(\mathbf{s}^{\mathbf{r}})$ is decoded from the spatiotemporal neuronal responses (Eqs. S22 and S23). And then we search the prior precision $\mathbf{L}$ from which the posterior is closet to the actual sample distribution $p(\mathbf{s}^{\mathbf{r}})$,

$$\hat{\mathbf{L}} = \arg\min_{\mathbf{L}} D_{KL}[p(\mathbf{s}|\mathbf{x})||p(\mathbf{s}^{\mathbf{r}})], \tag{S27}$$

where the parameter of posterior $p(\mathbf{s}|\mathbf{x}) = \mathcal{N}(\mathbf{s}|\boldsymbol{\mu}_{\mathbf{s}}, \mathbf{\Lambda}^{-1})$ is computed by,

$$\mathbf{\Omega} = \mathbf{\Lambda} + \mathbf{L}, \quad \boldsymbol{\mu}_{\mathbf{s}} = \mathbf{\Omega}^{-1}\mathbf{\Lambda}\mathbf{x}, \tag{S28}$$

and the likelihood precision $\mathbf{\Lambda}$ in above equation is decoded from the feedforward inputs (Eq. S26)

Then the off-diagonal elements of the estimated $\hat{\mathbf{L}}$ will be compared with the theoretical prediction (Eq. 17), and the results are shown in Fig. 3E and H.
